# Immortalization of human nasal and bronchial airway epithelial cells for genome editing applications

**DOI:** 10.1101/2025.11.30.691384

**Authors:** Emma J. van Grinsven, Georgia N. Ithakisiou, Pim Cleijpool, Bram M. Bosch, Marinos Tziouvelis, Gimano D. Amatngalim, Sam F. B. van Beuningen, Wilco Nijenhuis, Bahar Yetkin-Arik, Lukas C. Kapitein, Jeffrey M. Beekman, Anna Akhmanova

## Abstract

*In vitro* air-liquid interface culture of airway epithelial cells is used as a model system to study respiratory diseases. This culture system not only overcomes the need for animal models or continuous biopsies from individuals but also enables studies of pathophysiology associated with the disease in a patient background. Human airway basal cells serve as progenitor cells for a functional pseudostratified airway epithelium composed mainly of multiciliated and secretory cells. However, due to the limited ability of basal cells to proliferate and differentiate, the long-term use of primary material in culture is restricted. This challenges research that requires genome editing. Here, we describe airway stem cells from nasal and bronchial origin immortalized by hTERT overexpression followed by polyclonal expansion. We demonstrate that this diverse panel of cell lines shows differentiation patterns similar to primary stem cells and can be used for lentiviral and CRISPR/Cas9 genome editing. These cell lines and optimized protocols facilitate airway biology research and disease phenotyping.

## Introduction

The human airway epithelium lines the respiratory tract, providing protection against pathogens and particles, while supporting efficient gas exchange in the lower airways (Crystal et al., 2008; Knight and Holgate, 2003; Thompson et al., 1995). This epithelium is composed of several cell types, including basal cells that act as the proliferating and regenerating population of the epithelial sheet, goblet cells that produce mucus, and multiciliated cells that beat in a coordinated manner to remove mucus deposited by goblet cells from the distal to proximal airways (Bustamante-Marin and Ostrowski, 2017). For decades, air-liquid interface (ALI) culture models have been widely used to study differentiated primary human airway epithelial cells *in vitro* (Lee et al., 2022; Whitcutt et al., 1988). This is especially useful for studying epithelial defects in monogenic diseases, such as cystic fibrosis and primary ciliary dyskinesia (PCD) (Keegan and Brewington, 2021; Wallmeier et al., 2020). However, in contrast to commonly used cell lines, primary basal airway cells have limited proliferative capacity, and long-term airway culture is impossible (Bukowy-Bieryllo, 2021; Fulcher et al., 2009; Orr and Hynds, 2021; Widdicombe et al., 2005; Zabner et al., 2003). Furthermore, obtaining patient material is both time-consuming and costly, does not provide a scalable source of biological tissue and is subject to significant donor-to-donor variation, which can affect reproducibility and consistency in experimental outcomes. This limits the development of therapeutic strategies and studies regarding protein function using genetically modified airway epithelium, because genome manipulations often require multiple rounds of selection and thus long-term culturing (Bukowy-Bieryllo, 2021).

To address this problem, various strategies have been used to extend the time that airway epithelial cells can be maintained in culture. One approach involves technical advancements by optimizing culture conditions or by culturing airway cells in co-culture systems (Dreyer et al., 2024; Peters-Hall et al., 2020; Vaughan et al., 2006). In addition, genetic approaches have been employed to overcome replicative senescence, typically by introducing oncoproteins, such as the simian virus 40 large T antigen (SV40), human papillomavirus (HPV) E6 and E7 proteins or Bmi-1, or by manipulating pathways related to cell cycle regulation, for example, by expressing cyclin-dependent kinase (Cdk) 4 or by expressing human telomerase reverse transcriptase (hTERT), either alone or in combination with oncogenes (Cozens et al., 1994; Fulcher et al., 2009; Halldorsson et al., 2007; Lee et al., 2022; Lundberg et al., 2002; Piao et al., 2005; Ramirez et al., 2004; Reddel et al., 1988; Sato et al., 2020; Selo et al., 2021; Walters et al., 2013; Wang et al., 2019; Zabner et al., 2003). These immortalized airway cell lines have an extended life span and, to various degrees, retain characteristics of differentiated airway epithelium. For instance, although, 16HBE140 and BEAS-2B can be used in studies of drug absorption and transport, they do not form polarized epithelial sheets or differentiate into multiciliated cells (Cozens et al., 1994; Reddel et al., 1988). Other cell lines, such as VA10, BCi-NS1.1, huAEC and UNCN1T, support multiciliated cell differentiation, but often with low proportion of ciliated cells or reduced ciliary beat frequency compared to primary cells (Benediktsdottir et al., 2013; Kuek et al., 2018; Lundberg et al., 2002; Piao et al., 2005; Ramirez et al., 2004; Stewart et al., 2012; Vaughan et al., 2006; Walters et al., 2013; Wang et al., 2019). Consequently, extending the repertoire of immortalized lines is needed to exclude potential constraints associated with the use of cells that all share the same donor origin and to support the ongoing advances in respiratory disease research, gene-targeted therapies, and personalized medicine.

In the current study, we immortalized human airway epithelial basal cells using lentiviral expression of hTERT. We generated a panel of immortalized cell lines from bronchial and nasal origin and compared three nasal donors. The resulting cell lines retained the ability to differentiate into the major airway epithelial cell types and showed functional characteristics resembling those of healthy primary tissue material. Utilizing these immortalized cell lines, we showcased three genome editing strategies: lentiviral protein overexpression, gene knockout and knock-in. We expect that these immortalized cell lines in combination with the genome editing toolbox will help to extend our basic knowledge of airway processes and accelerate testing of new therapies.

## Results

### Immortalized airway basal cells exceed proliferation limit of primary cells and retain differentiation capacity

Primary cells were collected from human bronchial tissues and nasal brushings, and optimized culture conditions were used to selectively proliferate basal cells (Dreyer et al., 2024; Rodenburg et al., 2023b). Immortalized cell lines were generated by lentiviral expression of human telomerase reverse transcriptase (hTERT). Basal cells positive for integration were selected using hygromycin (Fig. 1A). Although serial dilution enabled isolation of individual clones in the past (Walters et al., 2013; Wang et al., 2019), cellular stress responses might be triggered due to low cell density. Here, we reasoned that bulk expansion of basal cells would naturally steer the population towards selection of the healthiest proliferating cells without clonal selection. To assess the variation and reproducibility of the immortalization strategy with hTERT alone, we cultured primary human epithelial cells obtained from two bronchial regions (HBEC1 and HBEC2) and from three nasal donors (HNEC1, HNEC2 and HNEC3), in parallel. While proliferation of primary HBECs and HNEC3 stalled around 40, 60 and 46 population doublings (PDs), respectively (Fig. 1B), cells from the same individuals expressing hTERT exceeded 78, 112 and 121 PDs, respectively, and could be cultured for over 300 days (Fig. 1C). Although the doubling time varied slightly from donor to donor, in all cell lines it remained constant over time. This demonstrates that hTERT overexpression alone is sufficient to generate immortalized bronchial cells (HBEC1i, HBEC2i) and nasal cells (HNEC1i, HNEC2i, HNEC3i), which exceed the proliferation limit of primary airway epithelium basal cells.

**Figure 1:**
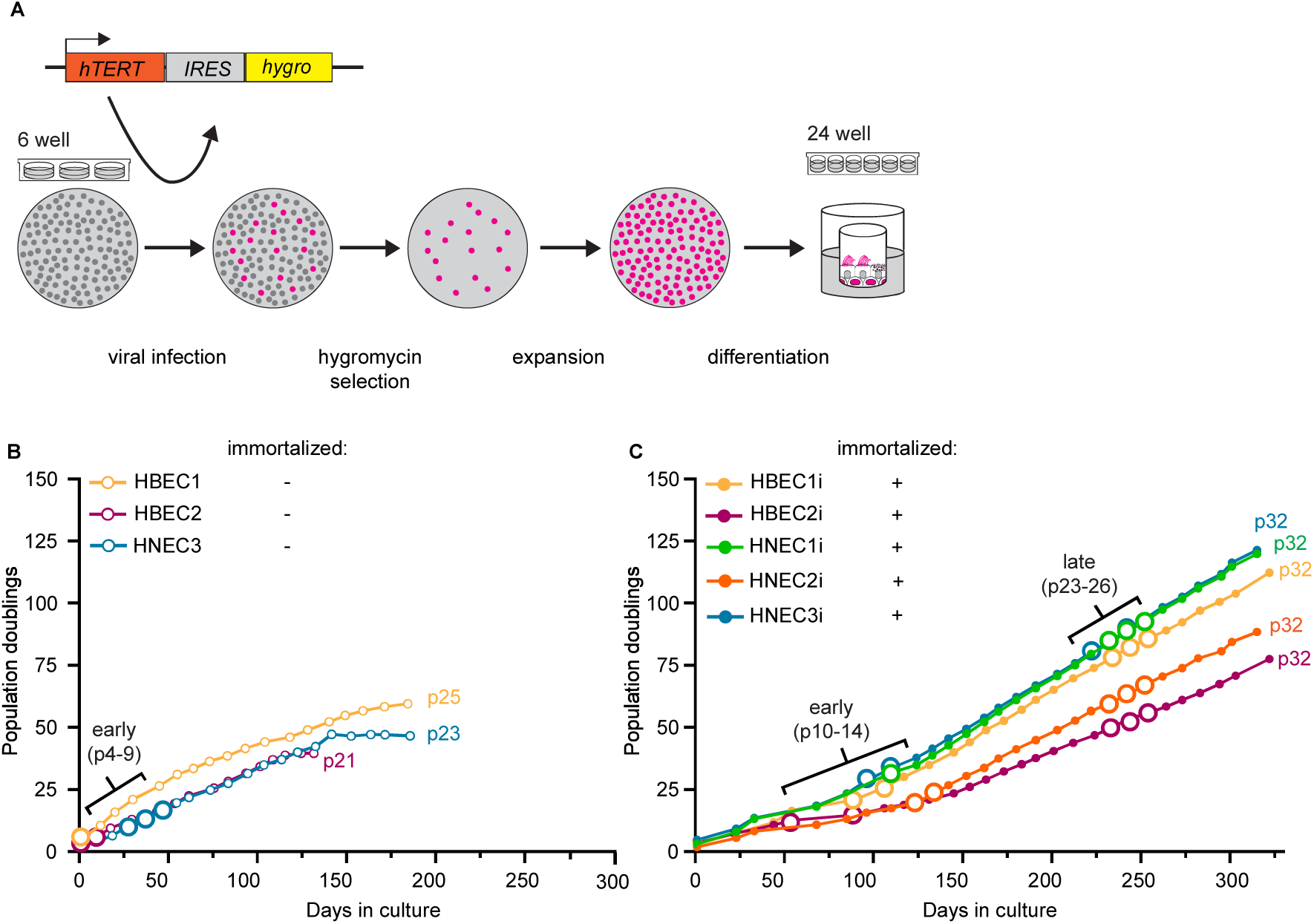
hTERT immortalized airway epithelial cells exceed proliferation limit. (A) Schematic overview of the workflow to generate immortalized airway cells. Primary basal cells were infected by lentiviral hTERT expression cassette; positive cells were selected using hygromycin, expanded and differentiated in ALI culture system. (B) Growth of primary bronchial (HBEC1 and HBEC2) and nasal airway (HNEC3) epithelial basal cells. Cell population doublings were plotted against time. Each point represents one passage (p) and the last passage number is indicated for each donor. The enlarged points represent early passages (p4-9) which were used for assays in Fig. 2. (C) Growth of immortalized (hTERT+) bronchial (HBEC1i and HBEC2i) and nasal airway (HNEC1i, HNEC2i, HNEC3i) epithelial basal cells. Cell population doublings were plotted against days in culture. Each point represents one passage, and the last passage number is indicated for each donor. The enlarged points represent early (p10-14) and late passages (p23-p26), which were used for the assays in Fig. 2.

To further characterize the immortalized cell lines, we compared their differentiation capacity at early (p10-14) and late (p23-26) passages to early passages (p4-9) of primary cells (HNEC3, HBEC1 and HBEC2) using immunofluorescence staining with antibodies against the major cell types present in the differentiated airway epithelium (Fig. 2A). Basal cells (P63) were present in all immortalized cell lines at both an early and late passage (Fig. 2A, B). Next, we assessed the presence of differentiated cell types (Fig. 2A-G). All primary cells and immortalized cell lines differentiated into β4 tubulin-positive multiciliated cells and MUC5AC-positive goblet cells (Fig. 2A, C, D). Immunostaining for CC16 and qPCR for SCGB1A1, both club cell markers, showed that these cells were rare or absent in all our primary and immortalized cell cultures, in line with their low numbers in the upper airway (Fig. 2E-G) (Blackburn et al., 2023). Because of our interest in multiciliated cell function, the culture medium was supplemented with the Notch inhibitor, DAPT, which promotes differentiation towards multiciliated cells at the expense of secretory cells, this expected shift in cell composition was clearly reflected in the cultures (Gerovac et al., 2014).

**Figure 2:**
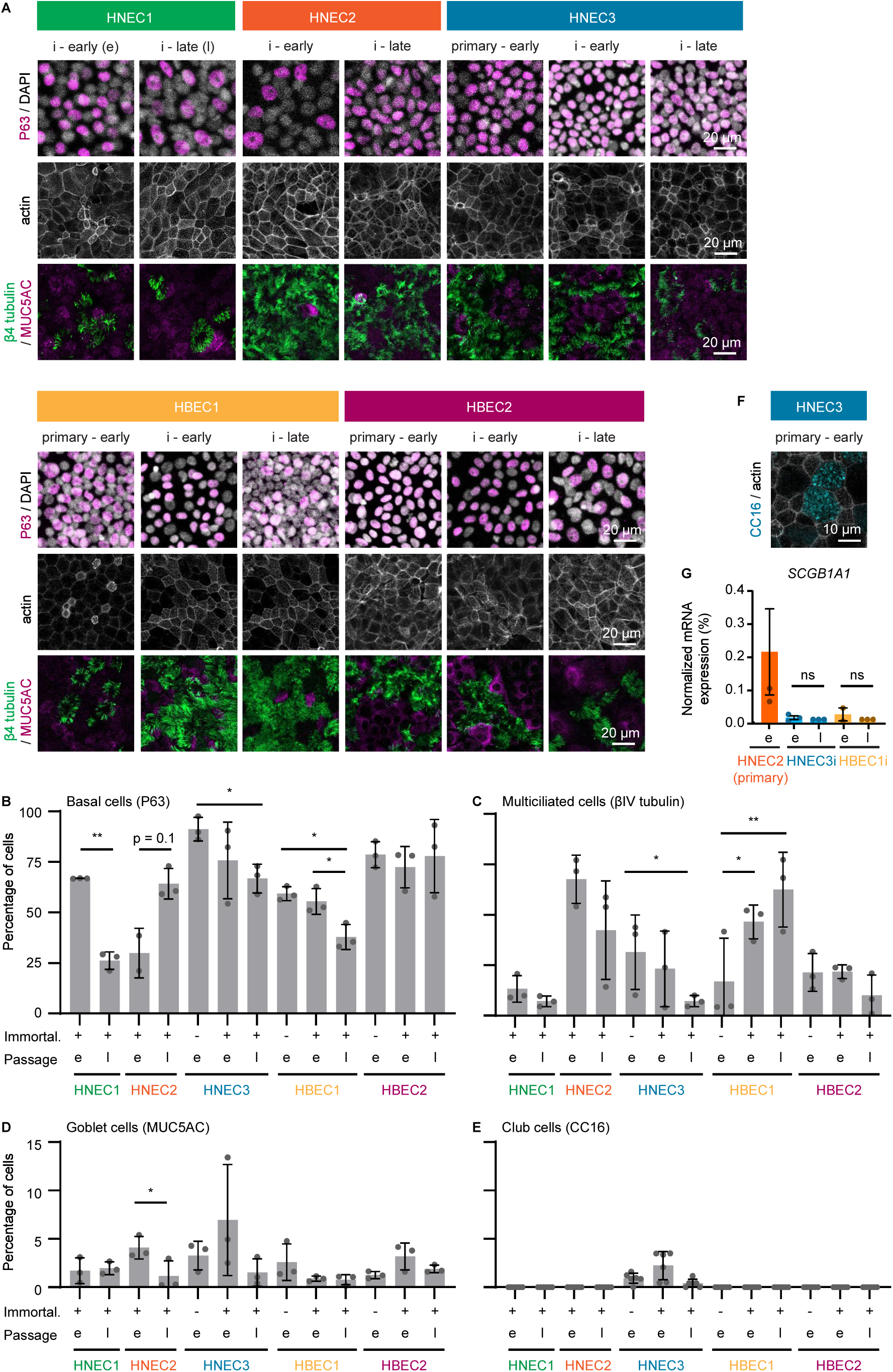
Immortalized airway cells retain the ability to differentiate into major airway cell types. (A) Immunofluorescence images of primary and immortalized (i) airway cells differentiated at early (e) and late (l) passages and stained for basal cells (P63), goblet cells (MUC5AC), multiciliated cells (β4 tubulin), nuclei (DAPI) and F-actin (phalloidin). P63 stained images were acquired on a Leica SP8, MUC5AC and β4 tubulin images were acquired on a Zeiss LSM980 Airyscan2. (B-E) Quantification of the percentage of basal cells (B), multiciliated cells (C), goblet cells (D) and club cells (E) in primary and immortalized airway cells at early (e) and late (l) passages. n, number of replicates: for B and E, n=3 with 1 field of views (FOVs) per replicate; for B and C, n=3 with 2 FOVs per replicate. Statistical tests were performed using an unpaired parametric T test with Welch’s correction. Data are shown as mean ± SD, and each point represents a replicate. (F) Immunofluorescence images of primary HNEC3 differentiated at an early passage and stained for club cells (CC16) and F-actin (phalloidin). Image was acquired on a Leica SP8. (G) Normalized mRNA expression levels to detect club cells (*SCGB1A1*) in indicated primary and immortalized airway cells differentiated at early (e) and late (l) passages. n, number of replicates: all conditions, n=3. Statistical tests were performed using an unpaired non-parametric T test. Data are shown as mean ± SEM, and each point represents a replicate.

Although all cultures contained the four major airway epithelial cell types, the relative proportions of these populations varied between donors and across passages. However, no consistent trend was observed, suggesting that these differences do not reflect donor-dependent variation but are more likely due to minor differences in culture conditions, as also indicated by variation between replicates. Together, these findings indicate that both bronchial and nasal cell lines immortalized by hTERT overexpression maintain the capacity to differentiate into mucociliary epithelium under optimized conditions.

### Immortalized airway cells retain functional characteristics of the differentiated epithelium

Next, we assessed several characteristic functions of airway epithelial cells differentiated from immortalized basal cells at late passage numbers (p24-26). First, ciliary beat frequency (CBF, Hz) was assessed by imaging cells using high-speed video microscopy. CBF of multiciliated cells differentiated from immortalized cell lines was between 6.5 Hz and 15.5 Hz and slightly lower than that of primary cells (between 7.7 Hz and 20.3 Hz) but remained constant throughout multiple days in ALI culture (Fig. 3A-C). Second, the barrier function of epithelial cells, which relies on the junctional integrity between cells, was measured by performing transepithelial electrical resistance (TEER) assays (Srinivasan et al., 2015) at different timepoints during ALI culture differentiation (Fig. 3D, E). Primary cells stabilized at approximately 1000 Ohms (Ω) per cm² at ALI day 7, whereas in immortalized cell lines differentiated at late passages (p24-p26), TEER reached ∼2000 Ω per cm² at ALI day 7 or 14. Lastly, an important function of airway epithelium is mucociliary clearance. In secretory cells, cystic fibrosis transmembrane conductance regulator (CFTR) protein modulates the transport of anions and fluid in and out of cells to control mucus consistency (Okuda et al., 2021). A forskolin-induced organoid swelling (FIS) assay can be used to examine CFTR efficiency and ion transport function (Amatngalim et al., 2022; Rodenburg et al., 2023a). We generated airway cystic organoids from one bronchial and one nasal immortalized cell line, HBEC1i and HNEC3i, respectively, at late passage number (p24 and p26) (Fig. 3F). FIS assay showed that organoids derived from both immortalized cell lines responded to forskolin by swelling approximately 2-fold in 1 hour, comparable to what has been published (Fig. 3F,G) (Amatngalim et al., 2022). This confirmed that the immortalized cell lines can be used to generate organoids suitable for measuring CFTR function with FIS assays.

**Figure 3:**
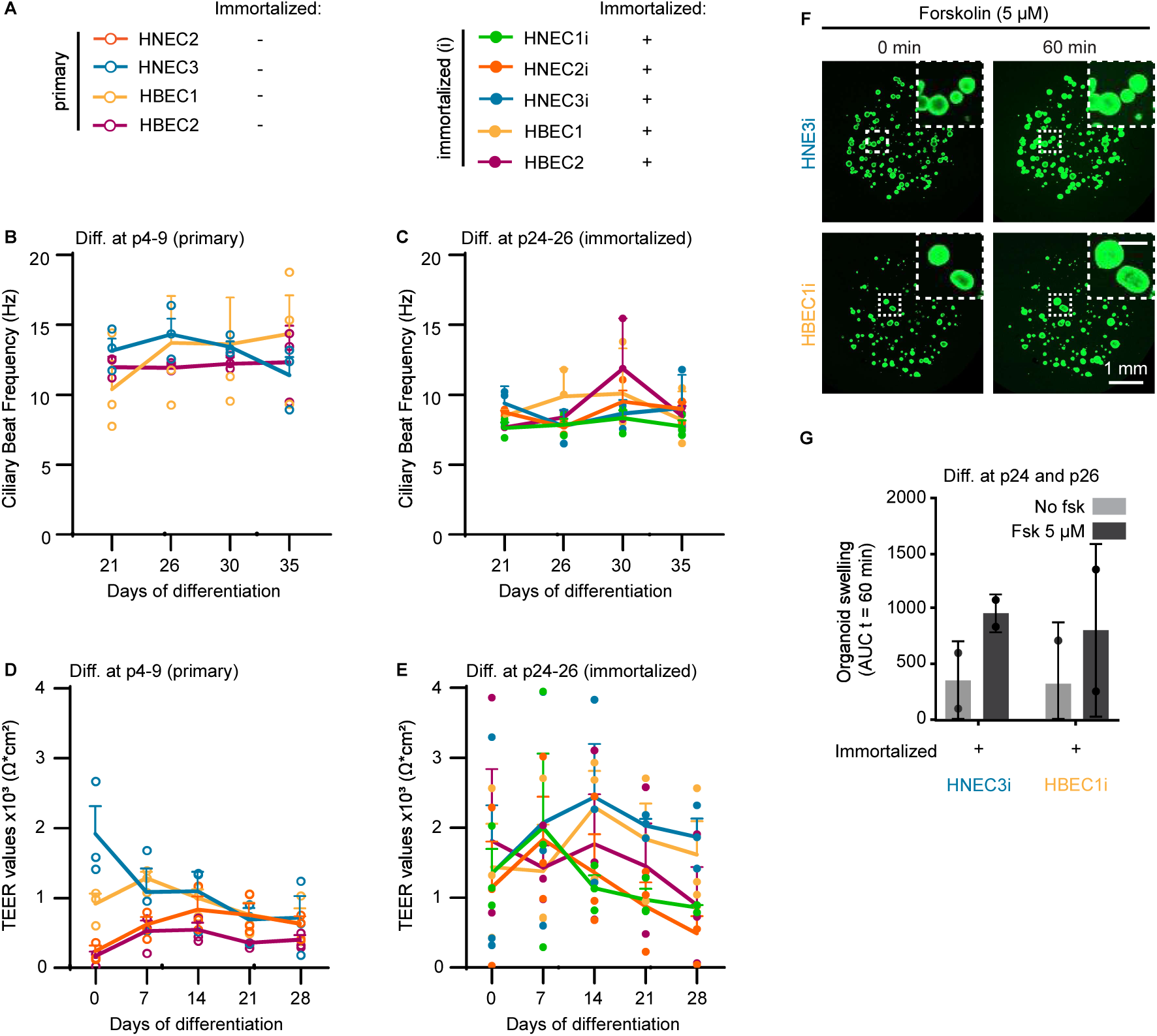
Functional assays with differentiated immortalized airway epithelial cells. (A) Legend indicating the colors used to depict data for primary donor cells or immortalized cell lines in (B-E). (B, C) Ciliary beat frequency (CBF) in hertz (Hz) of primary (p4-9) and immortalized airway epithelial cells differentiated at late passage number (p24-26). CBF was measured at several timepoints of differentiation. n, number of replicates: all conditions n=3. Data are shown as mean ± SEM, and each point represents a replicate. (D, E) Transepithelial electrical resistance (TEER) assay of primary (p4-9) and immortalized airway epithelial cells differentiated late passage number (p24-26), respectively. n, number of replicates: all conditions, n=3. Data are shown as mean ± SEM, and each point represents a replicate. (F) Representative images of forskolin-induced swelling determined with calcein green AM ester–stained immortalized HNEC3i and HBEC1i organoids, with images taken before the addition of 5 μM forskolin (0 min) and after stimulation (60 min). Insets show zooms of a few organoids. Scale bar in inset is 250 µm. Images were acquired on a Zeiss LSM800 (G) Quantification of area-under-the-curve (AUC) of HNEC3i or HBEC1i organoid swelling after 60 minutes in control conditions and after the addition of 5 μM forskolin. n, number of replicates: all conditions, n=2. Data are shown as mean ± SD, and each point represents a replicate.

Together, these assays demonstrate that immortalization does not affect airway epithelial functions such as ciliary beating, junctional integrity and ion and fluid secretion after long-term culturing. Therefore, they are suitable for modeling of diseases where ciliary activity or anion-dependent fluid secretion are affected.

### Lentiviral gene expression in immortalized HNEC

Having established immortalized airway epithelial cultures, we next used them for genetic modification. We first set out to visualize microtubules using overexpression of the well-characterized, fluorescently tagged β4B tubulin (encoded by *TUBB4B*, and recognized by β4 tubulin antibody) and microtubule-binding protein End Binding protein 3 (EB3) (Stepanova et al., 2003). Previous work on epithelial cells from mouse trachea and human lung tissue showed that lentiviral transduction is an effective strategy to express transgenes (Fulcher et al., 2009; Vladar and Brody, 2013). To this end, we infected HNEC3i basal cells with a lentiviral construct to express β4B tubulin with a N-terminal GFP tag and EB3 with a C-terminal tdTomato tag (Fig. 4A). Confocal microscopy of fixed basal cells showed that GFP-β4B tubulin and EB3-tdTomato decorated microtubules and EB3-tdTomato was enriched at microtubule plus ends as well, as expected (Fig. 4B). Furthermore, differentiation of this cell line confirmed that EB3-tdTomato specifically localized to ciliary tips, as described previously (Fig. 4C, D) (Schroder et al., 2011). This localization pattern was only visible in a subset of multiciliated cells due to the heterogeneity in EB3-tdTomato expression levels, likely caused by the variation in the number of lentiviral transgene inserts per cell (Paugh et al., 2021). This is a problem, especially when the localization of a protein of interest strongly depends on its expression levels, such as with EB3, which binds to microtubule tips at low expression but decorates the whole microtubules at higher expression levels (Fig. 4C). To overcome this heterogeneity, we selected basal cells with low (4.8% of total cells) and high (0.7% of total cells) EB3-tdTomato expression using fluorescence-activated cell sorting (FACS) of transduced basal cells (Fig. 4E). Live-cell imaging of differentiated HNEC3i expressing low levels of EB3-tdTomato with a Lattice Light-sheet microscope showed that EB3 specifically localizes to the tips of beating cilia in multiciliated cells (Fig. 4F). These optimization steps clearly illustrate the benefit of working with immortalized cells. Therefore, we conclude that lentiviral transduction is an effective approach to overexpress tagged proteins of interest for imaging of fixed and live airway epithelial cells generated from immortalized basal cells.

**Figure 4:**
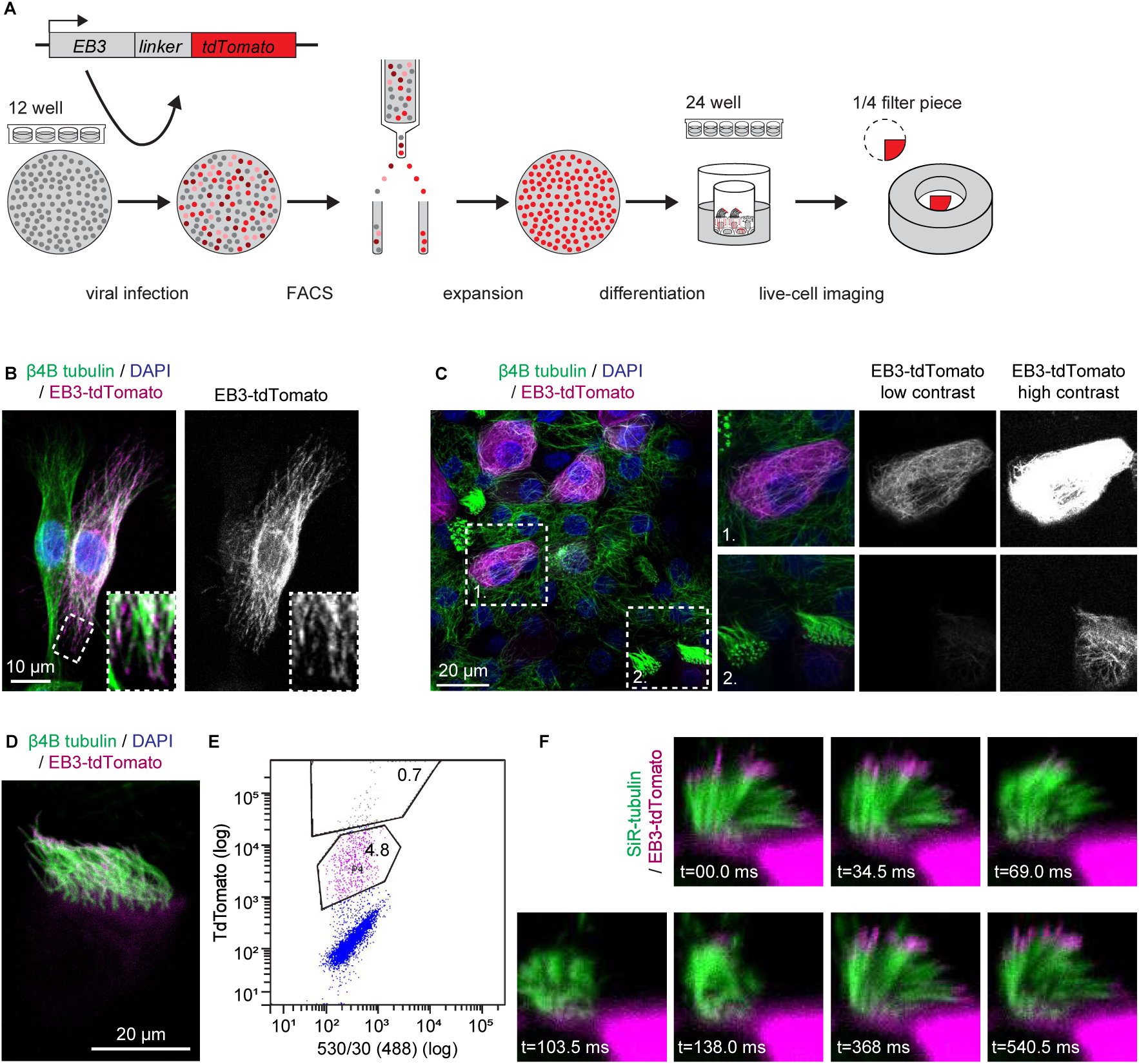
Lentiviral overexpression of EB3-tdTomato in immortalized HNEC3i. (A) Schematic overview of the workflow to generate HNEC3i expressing EB3-tdTomato. HNEC3i basal cells were infected with lentivirus containing EB3-tdTomato expression cassette; positive cells were enriched using FACS and expanded. HNEC3i EB3-tdTomato cells were differentiated in ALI culture. Quarter of a filter was removed, flipped and secured in an imaging ring for live-cell imaging. (B) Immunofluorescence image of HNEC3i basal cells expressing EB3-tdTomato (magenta) and β4B tubulin (green). Box indicates region of zoom. Image was acquired on a Zeiss LSM880 Fast Airyscan. (C) Immunofluorescence image of differentiated HNEC3i expressing EB3-tdTomato (magenta) and β4B tubulin (green). Boxes indicate zoomed regions: box 1 shows a non-ciliated cell with high EB3-tdTomato expression levels at low and high brightness and contrast setting, box 2 shows a ciliated cell with low EB3-tdTomato expression levels at low and high brightness and contrast setting. Images were acquired on a Zeiss LSM700. (D) Immunofluorescence image of cell shown in (C) box 2 in the cilia plane to illustrate the localization of EB3-tdTomato at ciliary tips. Images were acquired on a Zeiss LSM700. (E) Flow cytometry analysis of HNEC3i EB3-tdTomato. Cell selection based on TdTomato (Laser 561, Bandpass 582/15) signal intensity (y-axis) and green autofluorescence (Laser 488 nm, Bandpass 530/30) was used to improve cell separation (x-axis). Cells with low (P4) and high (P5) EB3-tdTomato expression levels are indicated. (F) Video frames of multiciliated cell visualized with live dye SiR-tubulin (green) and EB3-tdTomato (magenta) during one beat cycle imaged with a Lattice Light-sheet microscope.

### CRISPR genome-editing in immortalized HNECs: successful knockout and knock-in

To extend our genome editing toolbox, we explored the possibility of CRISPR-Cas9-mediated genome editing of immortalized HNECs. Previously, lentiviral transduction or electroporation were used to introduce the CRISPR-Cas9 machinery into airway epithelial basal cells (Chu et al., 2015; Vaidyanathan et al., 2022). In the former study, a single lentiviral vector was used to express sgRNA, Cas9 and a resistance gene to knock out the gene expressing a transmembrane glycoprotein, MUC18, used as a marker for cancer progression (Chu et al., 2015; Lehmann et al., 1989). In the latter study, a Cas9-sgRNA ribonucleic protein (RNP) complex was used together with a repair template divided over two adeno-associated viruses (AAVs) (Vaidyanathan et al., 2022). Here, we employed a CRISPR-Cas9 editing approach which surpasses the need for lentivirus, AAVs and multiple repair templates (Rodenburg et al., 2022). We focused on CCDC39, a protein required for the assembly of inner dynein arms and ciliary motility, the disruption of which is a major cause of PCD (Antony et al., 2013; Merveille et al., 2011).

To generate CCDC39 knockout cells, we electroporated three chemically synthesized sgRNAs targeting CCDC39 into HNEC1i, HNEC3i and HBEC1i basal cells in complex with Cas9 protein (Fig. 5A, B). Introduction of a Cas9/sgRNA complex has been shown to improve editing efficiency (Kim et al., 2014). Protein levels of CCDC39 in differentiated epithelium were analyzed using Western blotting (Fig. 5C). Comparison of CCDC39 signal to PCD patient material (HNEC_PCD1 and HNEC_PCD2) confirmed a successful knockout of CCDC39 in HNEC1i and HNEC3i and a strong decrease in protein levels in HBEC1i basal cells (Fig. 5C). Genomic sequencing and Inference of CRISPR Edits (ICE) analysis (Conant et al., 2022) showed that the knockout efficiency was 97%, 73% and 53% for HNEC1i, HNEC3i and HBEC1i, respectively. Those percentages are in line with the Western blot results, which show that HBEC1i is a heterogenous population with only partial knockout. Next, we analyzed ciliary beating in ALI differentiated knockout cell lines. Based on the live SiR-tubulin staining, the area occupied by multiciliated cells did not change upon CCDC39 knockout, but the percentage of multiciliated cells with beating cilia decreased compared to the wild type (Fig. 5D, E), whereas the CBF was unchanged (Fig. 5F). This confirms that electroporation of a sgRNA-Cas9 complex into immortalized cell lines is an effective strategy to generate knockouts, and that CCDC39 knockout recapitulates the phenotype observed in PCD patients (Abo et al., 2022; Antony et al., 2013).

**Figure 5:**
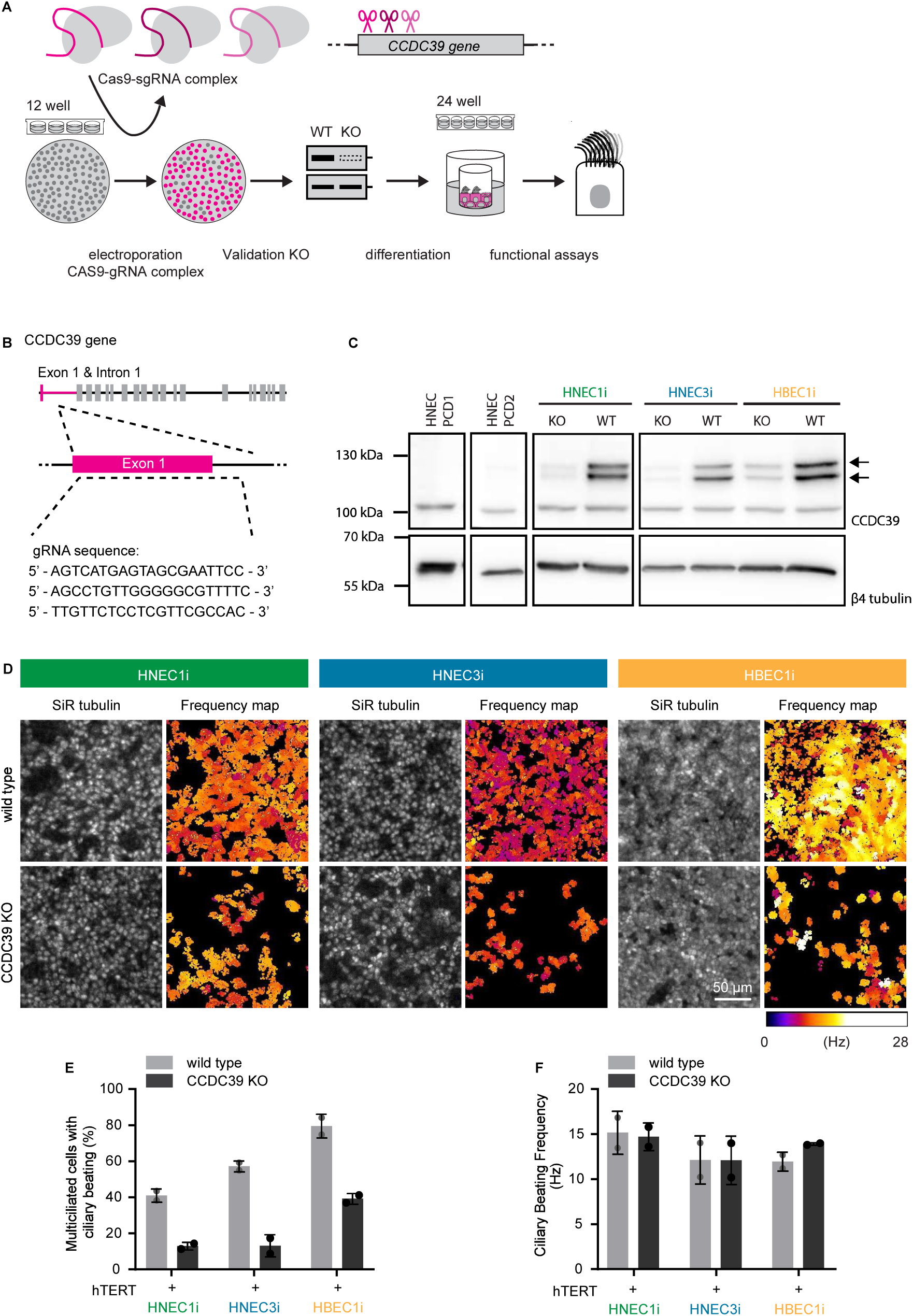
CRISPR-Cas9 knockouts of CCDC39 recapitulate PCD disease phenotype. (A) Schematic overview of CRISPR-Cas9 knockout strategy. Various immortalized airway basal cells were electroporated with sgRNA-Cas9 complex, differentiated in ALI culture, validated using Western blot and used for CBF assay. (B) Scheme of *CCDC39* and location of three sgRNAs. (C) Western blot illustrating the CCDC39 expression in PCD patient material (HNEC PCD1, HNEC PCD2) and wild type and knockout immortalized airway epithelial cells (HNEC1i, HNEC3i and HBEC1i). Arrows indicate CCDC39 protein. (D) Video frames of wild type and CCDC39 knockout (KO) cells HNEC1i, HNEC3i and HBEC1i incubated with live dye SiR-tubulin with corresponding CBF maps in hertz (Hz). (E,F) Area of multiciliated cells with beating cilia (E) and average CBF in Hz (F) in wild type and CCDC39 knockout (KO) cells HNEC1i, HNEC3i and HBEC1i. n, number of replicates: all conditions n=2. Data are shown as mean ± SD, and each point represents a replicate.

To test whether immortalized airway epithelial cells can also be used to obtain CRISPR-Cas9 mediated knock-ins, we generated knock-in cell lines where histone H2B (H2B) is labeled with mNeonGreen (mNG). HNEC3i basal cells were electroporated using two vectors, which contained mNeonGreen flanked by histone H2B-specific homology arms and H2B-specific gRNA and Cas9 (Fig. 6A). Ten days after electroporation, clones with green fluorescent nuclei could be readily observed, indicating successful integration of mNeonGreen into the histone H2B gene (Fig. 6B, left panel). Although knock-in efficiency was low, ∼0.05-0.1%, we could enrich positive cells using FACS to generate a 100% HNEC3i H2B-mNG positive cell line (Fig. 6B and 6C). The knock-in cell line retained the ability to differentiate (Fig. 6D). Notably, we could observe two types of cells, one with low and one with high fluorescent levels, suggesting a mixed population of heterozygous and homozygous histone H2B-mNG knock-in cells, respectively (Fig. 6E). We conclude that knock-in airway epithelial cell lines can be generated using electroporation of two vectors which together contain the repair template, gRNA and Cas9, and that positive cells can be enriched by using FACS.

**Figure 6:**
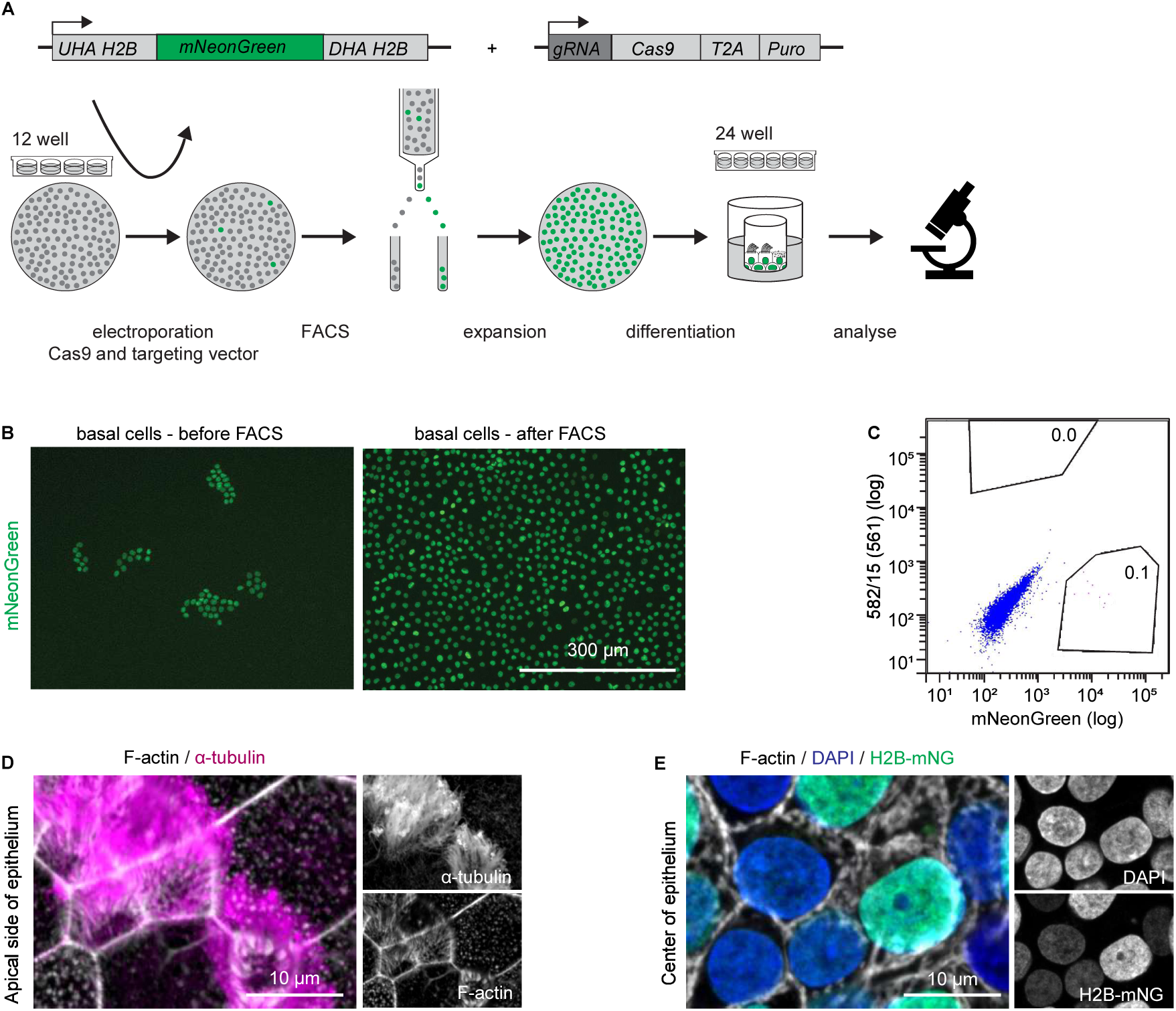
CRISPR-Cas9 histone H2B-mNeonGreen knock-in in immortalized HNEC3i. (A) Schematic overview of CRISPR-Cas9 knock-in strategy. HNEC3i basal cells were electroporated with two vectors containing mNeonGreen (mNG) flanked by up- and downstream homology arms (UHA and DHA) and histone H2B-specific guide RNA (gRNA), Cas9 and puromycin (puro) resistance gene. Positive cells were enriched by FACS, expanded and differentiated in ALI culture, and analyzed using immunofluorescence staining. (B) Representative EVOS microscope image of endogenous mNG signal in HNEC3i basal cells 10 days after electroporation (left) and after FACS (right). (C) Flow cytometry analysis of HNEC3i histone H2B-mNG. Cell selection based on mNG (Laser 488, Bandpass 530/30) signal intensity (x-axis) and red autofluorescence (Laser 561 nm, Bandpass 561/15) was used to improve cell separation (y-axis). Cells with high mNG signal were collected (P4). (D,E) Immunofluorescence image of sorted and ALI differentiated HNEC3i H2B-mNG cells stained for α-tubulin (magenta), F-actin (white), DAPI (blue) and mNG (histone H2B-mNG) fluorescence (green), showing cross sections at the apical side (D) or middle (E) of the epithelium. Images were acquired on a Zeiss LSM700.

## Discussion

In this study, primary human airway epithelial basal cells isolated from bronchial (HBEC) and nasal (HNEC) epithelium were successfully immortalized using lentiviral expression of hTERT to create the cell lines HBEC1i, HBEC2i, HNEC1i, HNEC2i and HNEC3i. The immortalized cell lines have been deposited in a biobank and are available to the research community on request. Immortalized cell lines retained the ability to differentiate into the major airway cell types, which displayed junctional integrity and ion transport function, and had motile cilia. Importantly, these airway characteristics remained stable in epithelial cells that were differentiated from immortalized basal cells of both early and late passages. Next, the validated, immortalized airway epithelial cell lines were used to optimize a genome editing toolbox for lentiviral overexpression of tagged proteins and CRISPR-Cas9 mediated genome editing.

Long-term airway epithelial cultures have been generated previously using hTERT expression in combination with the expression of oncoproteins or Cdk4, or co-culture with irradiated fibroblast feeder cells (reviewed in (Sato et al., 2020; Selo et al., 2021)). In the current study, similar to previous work (Piao et al., 2005; Walters et al., 2013; Wang et al., 2019), hTERT alone was sufficient to immortalize basal cells. Basal cells in this study and in the work from the Crystal lab were grown in optimized cell culture conditions, which by itself could be sufficient to reduce cell stress and promote the proliferative capacity of basal cells (Walters et al., 2013; Wang et al., 2019). This idea is in line with recent work suggesting that low oxygen, presence of the Rho-associated coiled coil protein kinase (ROCK) inhibitor and irradiated fibroblast feeder cells can extend the proliferation limit of primary bronchial airway epithelial cells to >200 PD (Peters-Hall et al., 2020).

Immortalization of human airway cells facilitates extended passaging and enables multiple rounds of selection, which are often essential for the validation of genome-edited cell lines. Here, we demonstrate with different genome editing workflows that a limited number of edited basal cells or a subset of labeled basal cells can be efficiently expanded to cryopreserve them, subsequently differentiate them and use them for functional assays. These techniques are instrumental to characterize the effects of complex, additive mutations, to conduct live-cell imaging studies or to visualize proteins in the absence of specific antibodies by knocking in different tags. Furthermore, they offer means to model rare, disease-associated mutations *in vitro*, particularly when patient-derived material is scarce or unavailable, whereas in the past, such studies were mostly conducted in genetically engineered mouse models (Gazdar et al., 2016). Moreover, donor heterogeneity in monogenic disease studies can be eliminated by employing isogenic lines or specifically addressed by inclusion of a panel of immortalized cell lines from multiple donors and origins. Lastly, these cell lines can provide ample material for large-scale assays for therapy development, including target identification and validation studies, and assessment of the effects of toxins or pharmacological agents on differentiated airway epithelial cells (Lee et al., 2021).

Taken together, we provided an easy-to-adopt genome editing toolbox which can help to address *in vitro* fundamental and disease-associated questions concerning airway epithelium without continuous dependency on donor material or animal models.

## Acknowledgements

This work was supported by the Netherlands Organization for Scientific Research (NWO) Gravitation programme IMAGINE! (project number 24.005.009) and the EindhovenWageningen-Utrecht Alliance (www.ewuu.nl) that supports the Centre for Living Technologies. We thank Didier Trono (EPFL, Switzerland) for providing pMD2.G and psPAX2 and Hugo Snippert (Princes Maxima Center for pediatric oncology, the Netherlands) for providing pSpCas9(BB)-2A-Puro with histone H2B-specific homology arms and the plasmid with H2B-specific gRNA and Cas9. We thank the Utrecht Platform for Organoid Technology (UPORT) for facilitating the ethical collection, biobanking, and distribution of human airway epithelial materials used in this study.

## Methods

### Isolation, expansion, and biobanking of primary airway epithelial cells

All individuals included in this study provided broad informed consent for the use of their samples in research. The study was reviewed and approved by the Biobank Research Ethics Committee (TCbio) of the University Medical Center Utrecht (protocol IDs: 16/586 and 22U-0079). The collection, storage, and distribution of airway epithelial materials were coordinated through the Utrecht Platform for Organoid Technology (UPORT) biobank, ensuring standardized procedures, ethical oversight and availability as resource material. Individuals characteristics (year of birth, sex, and relevant clinical history) are listed in Table 1. Primary nasal epithelial cells (HNEC) were isolated from healthy or PCD individuals through nasal brushing samples and cultured in nasal cell isolation medium (Table 1). Human bronchial epithelial cells (HBEC) were isolated either from non-tumorous resected airway tissue (HBEC1) or from bronchial biopsies of a patient with non-pulmonary cancer (HBEC2) using bronchial cell isolation medium. Bronchial cell isolation and expansion were performed as describe previously, and from this point onward cells were maintained at 37°C and 5% CO_2_ (Amatngalim et al., 2022; Dreyer et al., 2024). Briefly, after isolation, cells were seeded on a 6-well plate, which was pre-coated with collagen IV (50 µg/ml, Sigma Aldrich, Darmstadt, Germany) at 37°C and 5% CO_2_ for at least 1 h. After 7 days, the nasal or bronchial cell isolation media were replaced with nasal or bronchial cell expansion medium for nasal- and bronchial-derived cells, respectively (Table 1). The medium was refreshed every 2-3 days, and after reaching 80-90% confluency, the cells were passaged using TrypLE express enzyme (Thermo Fisher Scientific, MA, USA). At each passage, cells were counted and seeded at a density of 0.5 × 10⁶ cells per well of a collagen IV coated 6-well plate or cryopreserved using 500 µl CryoStor CS10 freezing medium (StemCell Technologies Inc., Vancouver, Canada) supplemented with 5 µM Y-27632 (Selleck Chemicals, TX, USA) at 0.5 × 10^6^ cells per vial. The proliferation rate of the cells was calculated by measuring population doublings (PDs) at each passage. PDs were defined as the log(Nf)-log(Ni)/log(2), where Nf is the final number of cells after expansion of each subculture and Ni is the initial number of cells that was plated.

### Differentiation of airway cells in air-liquid-interface (ALI) cultures

Primary and immortalized basal cells were grown and differentiated on 0.4 μm pore size polyester membrane Transwell inserts (Corning, NY, USA) coated with collagen I (PureCol Type I Bovine Collagen Solution; 30 µg/ml; Advanced Biomatrix, CA, USA) (Rodenburg et al., 2023a). Briefly, cells were seeded at a density of 0.2 × 10⁶ cells per insert in nasal or bronchial cell medium for nasal- or bronchial-derived cells, respectively. Cells were cultured with 200 µL medium on the apical side and 800 µL on the basolateral side, and the medium was refreshed every 2–3 days (5–7 days). Once confluent, the medium was changed to air-liquid interface (ALI)-differentiation medium (Table 1) supplemented with 500 nM A83-01; initially cells were cultured with 200 µL medium on the apical side for 5-7 days. Then, the apical volume was lowered to 45 µL (Dreyer et al., 2024). After 3-5 days, the apical 45 µL and basal 600 µL medium were replaced by ALI-differentiation medium supplemented with 20 µM DAPT (N-[N-(3,5-Difluorophenacetyl)-L-alanyl]-S-phenylglycine t-butyl ester) to promote differentiation towards ciliated cells (the experimental day 0 of differentiation). After ∼15 days, the apical medium was removed to create an ALI. For the knockout experiment, medium was supplemented with 5 µM DMH1 (Dreyer et al., 2024). During air-exposed differentiation, the medium was refreshed twice per week, and the apical side was washed once a week with PBS (Thermo Fisher Scientific, MA, USA). All cultured cells were regularly inspected using an EVOS M5000 microscope equipped with a 20x LWD PH, 0.45NA / 6.12 WD (AMEP4982) objective (images shown in Fig. 6B) or Leica DMi1 microscope equipped with a 20× HI PLAN I PH1 objective, 0.30 NA / X mm WD (456UE/03).

### Lentiviral production and infection

Lentiviruses were produced by a MaxPEI-based co-transfection of envelope vector pMD2.9 and packaging vector psPAX2. In brief, the supernatant of the packaging cells was harvested after 48 h and 72 h after MaxPEI-based transfection, filtered through a 0.45-μm filter and incubated with a polyethylene glycol (PEG) 6000-based virus precipitation buffer at 4°C overnight. Virus suspension was then centrifuged at 4°C and 1500x g for 30 min, the supernatant was removed, and the lentivirus was resuspended in PBS.

Basal cells were seeded to achieve 50% confluency in a 12-well plate at the day of infection. Prior to infection, basal cells were incubated with 5 μg/mL Polybrene (Millipore) for 1 h. 2.5 μL lentivirus was added in a dropwise manner. After infection with lentivirus containing hTERT (Hayer et al., 2016), positive cells were selected using 100 μg/mL G418 (Geneticin G418, InvivoGen) until control basal cells died.

### CRISPR-Cas9 genome editing

Generation of CRISPR-Cas9 cell lines was achieved by electroporation of immortalized cells with either ribonucleoprotein (RNP) complex or two DNA vectors (Rodenburg et al., 2022). For CCDC39 knockout, RNP complexes were prepared by mixing recombinant 2NLS-Cas9 nuclease (20 μM, 2.5 μL), multi-guide RNA (mix of three guide RNA’s, see Table 1) (30 μM, 8.3 μL) (chemically synthesized by Synthego) against CCDC39 and 14.2 μL OptiMEM supplemented with 10 μM Y-27632 and incubated at room temperature for 10 min. Next, 1.0 × 10^6^ cells were diluted in 75 μL optiMEM supplemented with 10 μM Y-27632 and added to the RNP complexes. Electroporation was carried out using the NEPA21 system according to the previously published protocol (Fujii et al., 2015). Following electroporation, cells were plated in 6-well plates containing nasal or bronchial cell medium.

For H2B-mNG knock-in, 2.5 μg vector with histone H2B specific sgRNA and Cas9 (Ran et al., 2013) and 7.5 μg vector with mNeonGreen flanked by histone H2B homology arms (gift from H. Snippert, Princes Maxima Center for pediatric oncology, the Netherlands)(Bollen et al., 2022) were premixed and stored on ice. Basal cells were trypsinized, and 0.5 × 10^6^ cells were resuspended in Neurobasal medium (Gibco) supplemented with 1% B-27 supplement, 0.5 mM glutamine, 15.6 μM glutamate and 1% penicillin/streptomycin. Basal cells were mixed with premixed DNA vectors (and protein), collected in 10 μl electroporation tips and electroporated 2 times with a 20 ms 1500 Volt pulse using a Neon electroporation system (Invitrogen™ MPK5000). Immediately after electroporation, basal cells were plated in a 12-well plate and cultured for 24 h in the absence of antibiotics to promote cell survival.

### Inference of CRISPR Edits (ICE) analysis

Quantification of the knockout efficiency of HNEC cultures was performed by first isolating the DNA from 0.2 × 10^6^ cells using the Quick-DNA Microprep Kit (Zymo Research, CA, USA) following the manufacturer’s guidelines. Target regions were amplified by PCR using GoTaq G2 Flexi DNA polymerase and specific gene primers (Table 1). PCR products were separated by size on a 1% TAE-agarose gel. DNA fragments were excised and purified with the QIAquick Gel Extraction Kit (QIAGEN N.V., Germany). Purified DNA samples, along with sequencing primers (1:1), specifically designed to target the edited regions, were sent to Macrogen for Sanger sequencing. Knockout efficiency and specific deletions were analyzed using the ICE analysis tool (https://ice.synthego.com/#/). The calculated CCDC39 KO efficiency was 97 % for HNEC1i, 73 % for HNEC3i, and 53 % for HBEC1i.

### FACS enrichment of HNEC

Fluorescence-activated cell sorting (FACS) was used to select positive H2B-mNeonGreen and EB3-tdTomato cells. Basal cells were trypsinized, collected in the wash buffer and centrifuged at 400 x g for 5 min (Table 1). Cells were then resuspended in a small volume of 0.5% FBS in PBS at a density not exceeding 20.0 × 10^6^ cells / ml. Basal cells were sorted on a BD FACSAria Fusion Sorter (BD Bioscience) equipped with 4 lasers (blue (488 nm), yellow/green (561 nm), red (640 nm) and violet (405 nm)), a 100-μm nozzle size and 20 ψ pressure and collected in a 15 mL tube containing bronchial cell expansion medium. Collected basal cells were centrifuged for 5 min at 400 x g before being plated in a 24-well or 12-well plate, depending on the yield.

### Immunofluorescence cell staining

Cells were fixed in pre-warmed 4% paraformaldehyde (PFA) at 37°C for 15 min or in ice-cold MeOH at -20°C for 15 min. Prior to fixation of differentiated airway epithelial cells, the apical cell side was washed with 125 μl pre-warmed PBS at 37°C for 5 min to remove mucus. This step was left out in case basal cells were stained. Fixative was removed and sample was washed three times with PBS for 10 min. For immunofluorescence staining, the filter was carefully sectioned from the Transwell using a scalpel, and only a quarter piece was used to continue. For PFA fixed samples, cells were permeabilized in 0.5% Triton X-100 for 1h. Sample was then blocked in 3% BSA in PBS (blocking buffer) for at least 1 h at room temperature before incubation with primary antibodies diluted in the blocking buffer (Table 1). The samples were incubated with primary antibodies in a 0.5 μl Eppendorf tube on an orbital shaker for 4 h at room temperature or at 4°C overnight. Primary antibodies were removed using 3 washes in PBS for 10 min, and the sample was incubated with secondary antibodies diluted in blocking buffer for 4 h at room temperature. Secondary antibodies were removed by 3 10 min washing steps with PBS, dehydrated in 70% EtOH followed by 100% EtOH, airdried and mounted in Prolong Diamond or Vectashield using 13 mm thickness 1.5 coverslips.

Fixed samples were imaged on Zeiss LSM980 Airyscan2 microscope equipped 405 nm, 445 nm 488 nm 561 nm and 639 nm lasers, Axiocam 305 mono R2, 5.07 megapixel, 2464×2056, SONY IMX264 CMOS sensor with a PL APO 20x/0.8 Air WD=0.55mm; Leica TCS SP8 STED 3X microscope with continuous 405 nm and pulsed (80 MHz) white-light lasers, PMT and HyD detectors and spectroscopic detection with a PL APO 20x/0.75 IMM CORR CS2 objective; Zeiss LSM700 microscope equipped with 405 nm, 488 nm, 555 nm and 633 nm lasers and Plan-Apochromat 63x/1.40 Oil DIC objective. Zeiss LSM880 Fast Airyscan microscope equipped with 405 nm, Argon Multiline, 561 and 633 nm lasers, Alpha Plan-APO 63x/1.2 Oil DIC objective with PMT detectors.

### Quantification of cell types

Quantification of airway cell types was performed by staining for basal cells (P63), club cells (CC16) or goblet (MUC5AC) and multiciliated cells (β4 tubulin) together. To quantify the total number of cells in a field of view (FOV), the nuclei (DAPI) were automatically counted using FIJI plugin StarDist 2D on a MAX projection of the Z-slices containing the airway epithelial cell layer. The number of basal cells was also counted automatically using StarDist 2D. To visualize club cells, we acquired smaller regions at a higher resolution and counted their prevalence manually. Club cells were only observed in HNEC3 cells. Multiciliated cells often appeared in patches, and their cilia splayed onto neighboring cells. To accurately count the number of ciliated cells, we therefore used phalloidin, an actin marker with increased apical signal intensity in multiciliated cells. First, to correct for sample tilt, the large image was divided in 16 quadrants. For each quadrant, the most in focus patches were identified using Tenengrad focus (Tenenbaum, 1970). The five best-focused patches were selected, and the patch closest to the apical plane was max projected with its adjacent patches (z-1, z, z+1) (https://github.com/Living-Technologies/immortalized_lines_analysis). We used Cellpose3 to segment the phalloidin signal (Stringer and Pachitariu, 2025). Regions of multiciliated cells were annotated using a FIJI segment-anything-models 2 (SAM2) (Tiny model) plugin. The number of cells in each region was then determined using the phalloidin based cell outlines. Their position was validated using the β4 tubulin signal. Subsequently, the number of annotated multiciliated cells was counted using the previously generated phalloidin based cell outlines. To count the number of goblet cells, we first max projected the Z slices of the MUC5AC channel. We then measured the signal in this projection for every pixel in the manually annotated multiciliated cell regions and used these to calculate a mean (MUC5AC_background_mean) and standard deviation (MUC5AC_background_stdev) background signal. Every cell segmented by Cellpose was next analyzed, and if the mean value of the corresponding pixels exceeded our threshold (1*MUC5AC_background_mean + 2 *MUC5AC_background_ stdev), we assigned it as a goblet cell.

Per cell line and early or late passage number, two fields of view (FOVs) per replicate (n=3) were analyzed. Field of view areas were 0.207, 0.016, 0.180 and 0.180 mm² for basal cells, club cells, goblet cells and multiciliated cells, respectively.

### Lattice Light-sheet live-cell imaging

To perform live-cell imaging of differentiated airway epithelial cells, a 35 mm Attofluor Cell Chamber (Invitrogen) with a plasma cleaned 25 mm coverslip (Harrick Plasma Cleaner PDC-002) was assembled, and 30 μL of 10 μM 15 μm clear, silica particles (Kisker Biotech) in ALI-differentiation medium was added as a droplet to the center of the coverslip. Then, a quarter filter piece was carefully sectioned out of a Transwell using a scalpel and placed onto a cleaned and charged 18 mm coverslip (Harrick Plasma Cleaner PDC-002) with the basal layer towards to coverslip using tweezers. The 18 mm coverslip was then immediately inverted and carefully placed onto 30 μL droplet containing diluted beads. To this, ∼1 mL of ALI-differentiation medium was added before the chamber was closed using a second 25 mm coverslip. This setup ensures the autofluorescence of the filter does not interfere with the imaging. In addition, the beads act as spacers to prevent airway epithelial cells from being squashed between the coverslips. Lastly, the sealed environment and surplus of ALI-differentiation medium promote cell survival.

Live-cell imaging was performed on a Zeiss Lattice Lightsheet 7 microscope equipped with 488 nm, 561 nm and 640 nm lasers, a 13.3x/0.44 excitation objective and a 44.83x/1 observation objective; images were projected to the pco.edge4.2 (Excelitas PCO) camera with a final pixel size of 145 nm. For illumination we used Sinc3 beams with a 30 mm × 1000 mm lightsheet.

### Western blotting

CCDC39 was detected by Western blotting. Differentiated cells were trypsinized and lysed in Laemmli buffer (120 mM Tris-HCl pH 6.8, 20 % glycerol, 4 % sodium dodecyl sulfate (SDS), protease inhibitor (Fisher Emergo B.V.)), and the lysates were homogenized using a QIAshredder (QIAGEN N.V., Germany) column to reduce viscosity. Protein concentration was measured using the Pierce BCA Protein Assay Kit (Thermo Fisher Scientific, MA, USA) according to the manufacturer’s instructions. Optical density was recorded at 562 nm using a multimode plate reader, and protein concentrations were calculated based on a standard curve. For each sample, 15 μg of protein diluted in Laemmli buffer and 20x sample buffer (Laemmli buffer, 0.1 % bromophenol blue, 5 % β-mercaptoethanol) for a total volume of 45 μL as well as PageRuler Prestained Protein Ladder, 10 to 180 kDa (Thermo Fisher Scientific, MA, USA) were loaded on 7.5% SDS polyacrylamide gel for separation. Proteins were transferred to a PVDF membrane (Immobilon FL; Sigma-Aldrich, MO, USA) by overnight electrophoresis at 30 Volt in a cold room. Afterwards, the membrane was blocked at room temperature using 5% (w/v) non-fat milk powder (Campina, Amersfoort, Netherlands) in Tris-buffer saline with 0.05% Tween-20 (TBS-T) for 1 h. Rabbit primary antibodies against CCDC39 and β4 tubulin were diluted in 0.5% (w/v) non-fat milk powder in TBS-T and incubated for 3 hours at room temperature on a roller mixer. Following washing with TBS-T, the membrane was incubated with swine anti-rabbit-HRP secondary antibody diluted in 0.5% (w/v) non-fat milk powder/TBS-T and incubated at room temperature on a roller bank for 1 hour maximum. Chemiluminescence detection was performed using SuperSignal West Pico PLUS Chemiluminescent Substrate (Thermo Fisher Scientific, MA, USA) for β4 tubulin, and SuperSignal West Femto Extended Duration Substrate (Thermo Fisher Scientific, MA, USA) for CCDC39. Blots were visualized using the Chemidoc Touch Imaging system (Bio-Rad Laboratories, Inc., CA, USA).

### High speed video microscopy and quantification of ciliary beat frequency

The movement of the cilia in ALI cultures was analyzed by measuring the Ciliary Beating Frequency (CBF). ALI-differentiated HNECs were imaged on the Zeiss Celldiscoverer 7 microscope equipped with widefield LEDs 385 nm, 420 nm, 470 nm, 520 nm, 567 nm, 590 nm, 625 nm lasers, WF emission filters QBP 425 / 30+514 / 30+592 / 25+709 / 100 and TBP 467 / 24+555 / 25+687 / 145 and Axiocam 712 mono, and Plan-Apochromat 5x/0.35 WD 5.10 (dry) objective with magnification options 2.5×/0.12, 5x/0.25, 10x/0.35, utilizing the environmental control (O_2_ Module S, and TempModule S, Zeiss) to maintain cultures at 37°C and 5% CO_2_ during the measurement. The presence of cilia in the analyzed videos was verified using SiR-tubulin (Spirochrome, Thurgau, Swizerland). The stain was diluted in ALI-differentiation medium (1:5000) and added on the apical side of the Transwell inserts and incubated at 37°C and 5% CO_2_ for 4 h. For CBF assessment, videos were recorded with high-speed video microscopy (HSVM) at 4 random positions per culture at 216 frames per second (fps) for a total of 512 cycles, using a 5x magnification objective.

The recorded videos were analyzed using FIJI Correlesence v.0.0.6 plugin (“Temporal ICS” command) (https://github.com/ekatrukha/Correlescence), and the output of CBF analysis gave a frequency heatmap for each position from which both the CBF (Hz) and beating area (%) could be determined. Briefly, the mean intensity image was subtracted from each video stack to eliminate the static component. A normalized autocorrelation function was then calculated for each pixel over varying time delays. The oscillation period was determined by identifying the first maximum of the autocorrelation function, accepting values only if the peak exceeded a 0.2 threshold. The resulting frequency data were rendered as a heatmap image, with pixel intensity corresponding to frequency (Hz). Frequencies below 0.1 Hz and above 28 Hz were excluded. To reduce background noise, pixel clusters smaller than 25 pixels were removed. The final output provided both the mean CBF (Hz) and the mean of the beating area (%) for each analyzed position.

### Transepithelial electrical resistance (TEER) measurements

The integrity and permeability of the differentiated airway epithelium was evaluated by TEER measurements that were performed weekly throughout the differentiation period. Cells were apically washed and incubated with 150 µl PBS at 37°C and 5% CO₂ for 15 minutes. TEER was subsequently measured on the apical side using an EVOM2 voltmeter (World Precision Instruments) equipped with STX2 chopstick electrodes in the presence of 250 µl PBS. Transepithelial resistance (Ω·cm²) was calculated by subtracting the resistance of a blank, empty insert filled with 250 μl PBS, from the average value of two resistance readings. Finally, this value was multiplied by the surface area of the monolayer (∼0.33 cm² for each insert).

### RNA extraction, cDNA synthesis, and quantitative real-time PCR (qPCR)

Total RNA was extracted from differentiated ALI cultures using the NucleoSpin® RNA kit (MACHEREY-NAGEL, Duren, Germany); RNA concentration and purity were assessed using a NanoDrop spectrophotometer (Nanodrop, ND1000) and cDNA was synthesized using the iScript cDNA synthesis kit (Bio-Rad, Hercules, CA, USA) using the manufacturer’s protocols. cDNA at a final concentration of 500 ng·µl^−1^ was subjected to a two-step quantitative real-time PCR (qPCR) SYBR Green reaction on a CFX-384 real-time detection system (Bio-Rad, USA) using cell-type-specific primers (Table 1). qRT-PCR reaction was initiated with a 3-minute denaturation step at 95°C, followed by 40 cycles of 10 seconds at 95°C and 30 seconds at the annealing temperature at 63°C. For relative gene expression analysis, Ct values were normalized against housekeeping genes *ATP5B* and *RPL13A*, chosen for their stable expression in airway epithelial cells under varying experimental conditions, using the 2-ΔΔCT method (Livak and Schmittgen, 2001). Melt peaks were analyzed to confirm amplification of a single product. Three biological replicate experiments were performed with two technical replicates per condition.

### Forskolin-Induced Swelling (FIS) assay

The baseline function of the CFTR channel was measured using the FIS assay following a previously established protocol (Amatngalim et al., 2022; Rodenburg et al., 2023b; Vonk et al., 2020). Briefly, airway organoids derived from differentiated ALI cultures were plated in a 96-well plate and were stained for 30 minutes with calcein green AM (3 μΜ) (Invitrogen, MA, USA) diluted in DMSO. Following this incubation, organoids were exposed to 5 µM forskolin (Sigma-Aldrich, MA, USA). Images were acquired using a Zeiss LSM800 microscope equipped with 405 nm, 488 nm, 561 nm, and 640 nm lasers and 2.5/0.12 Air objective FWD=8.7mm to detect the calcein green signal, under controlled environmental conditions of 37 °C, with 95 % O₂ and 5% CO₂ (O_2_ Module S, and TempModule S, Zeiss). The swelling response was quantified by measuring the increase in the total area of calcein green AM-stained organoids at 10-minute intervals over 60 minutes.

### Statistical Analysis

Statistical analyses in Fig. 2 and 4 were performed in using Prism 8.4.3. and an unpaired, parametric t test with Welch’s correction, with * indicating p < 0.005 and ** indicating p< 0.01.

## Supplementary Data

**Supplementary Table 1:**
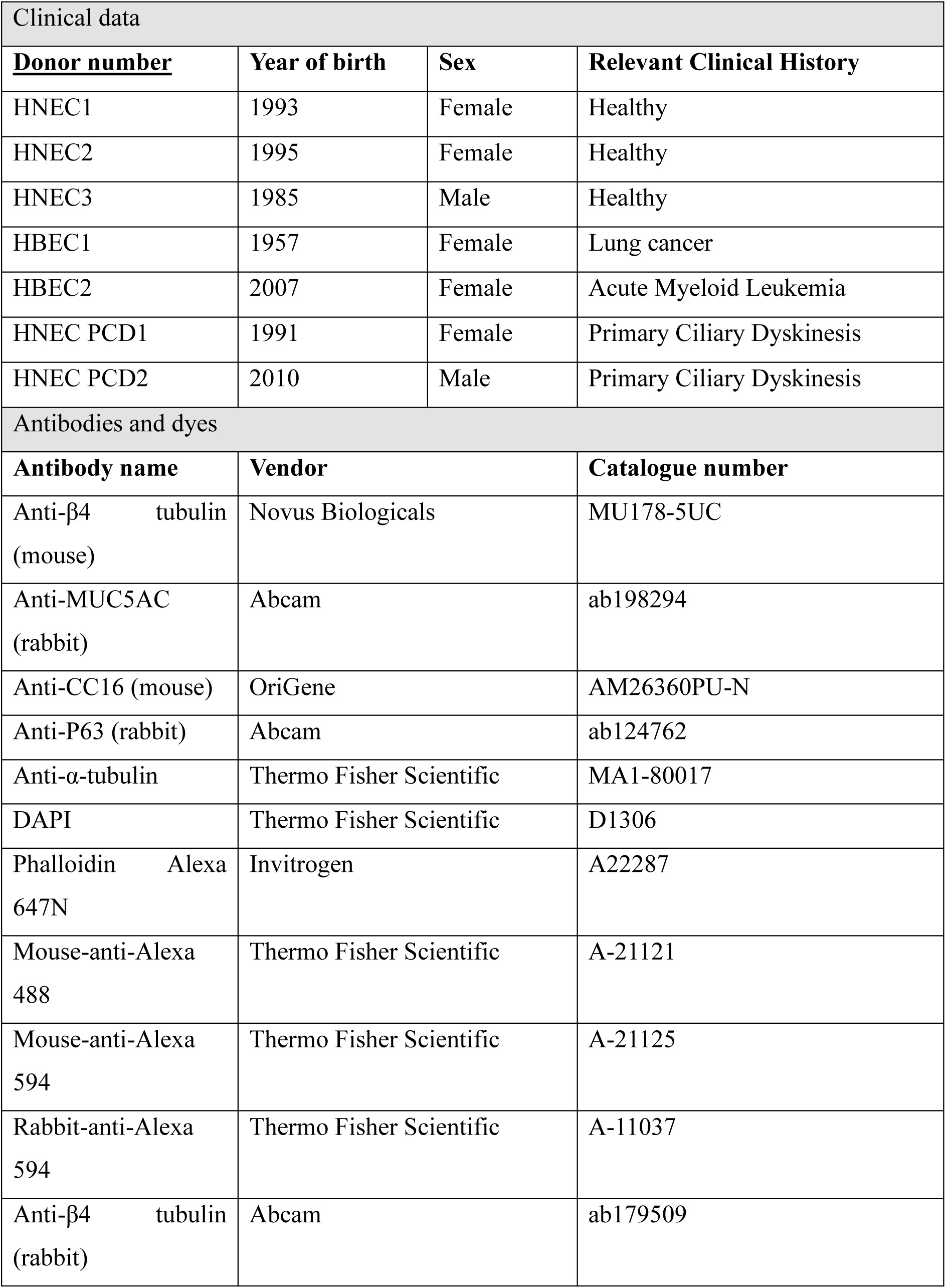
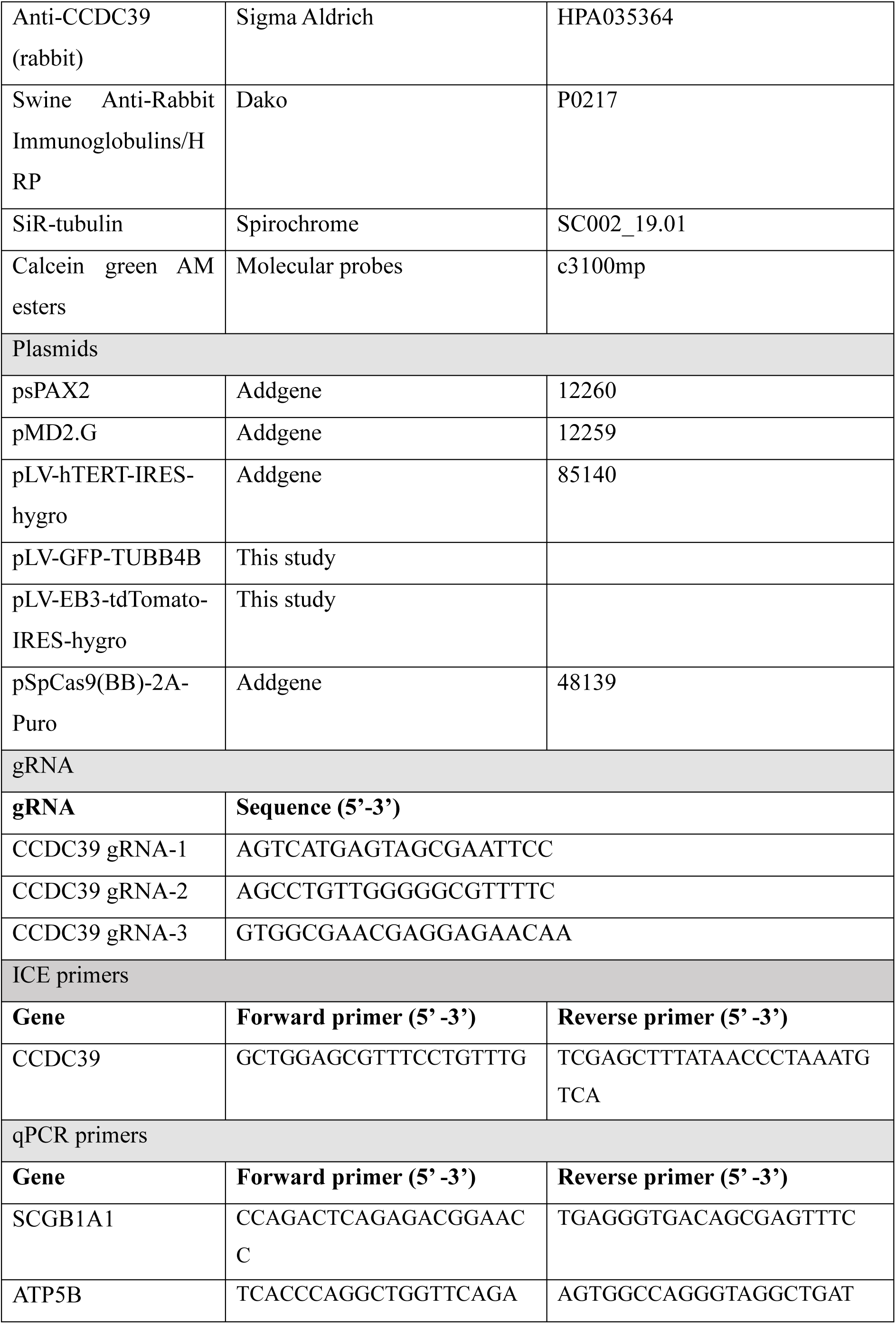
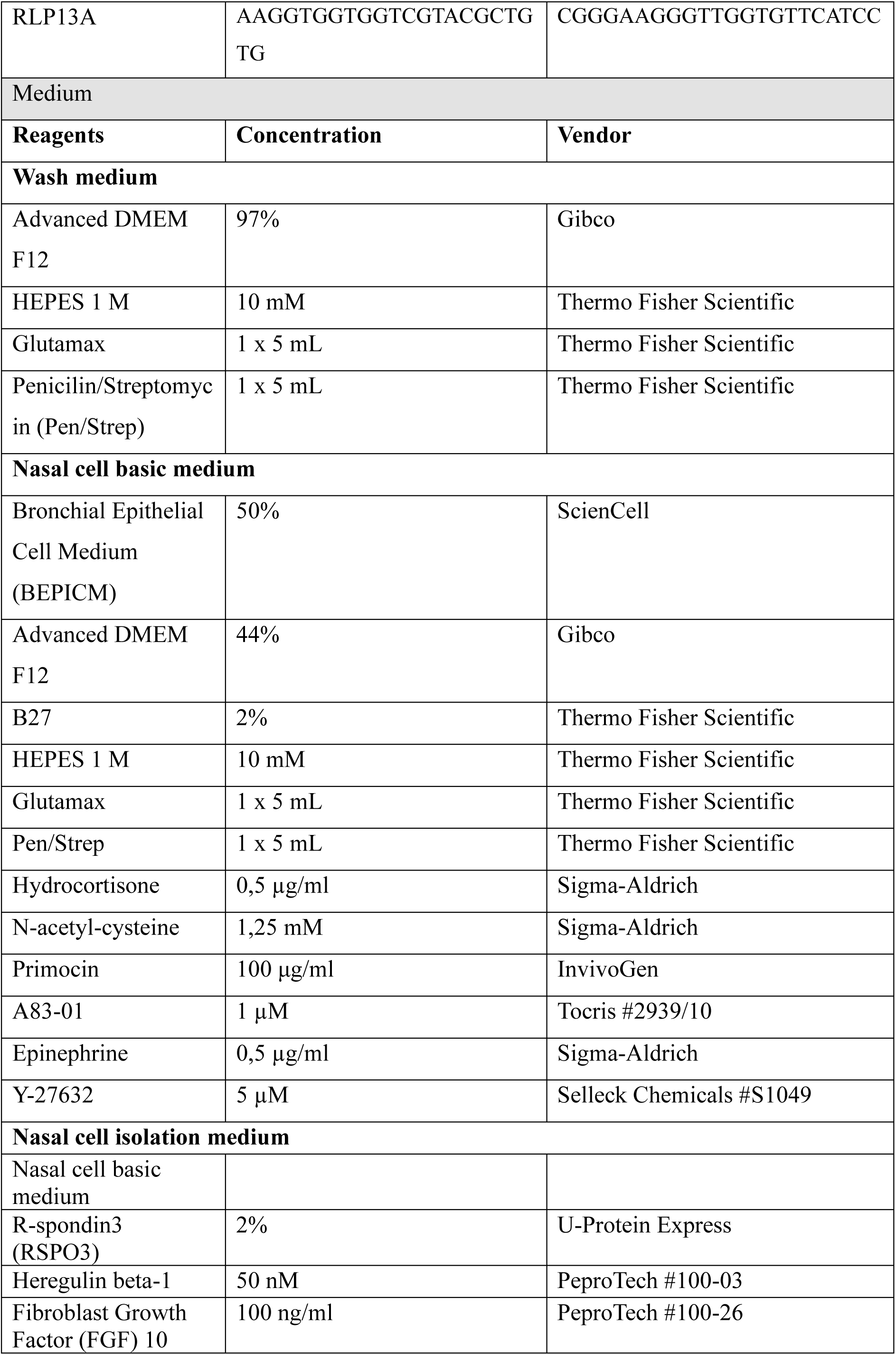
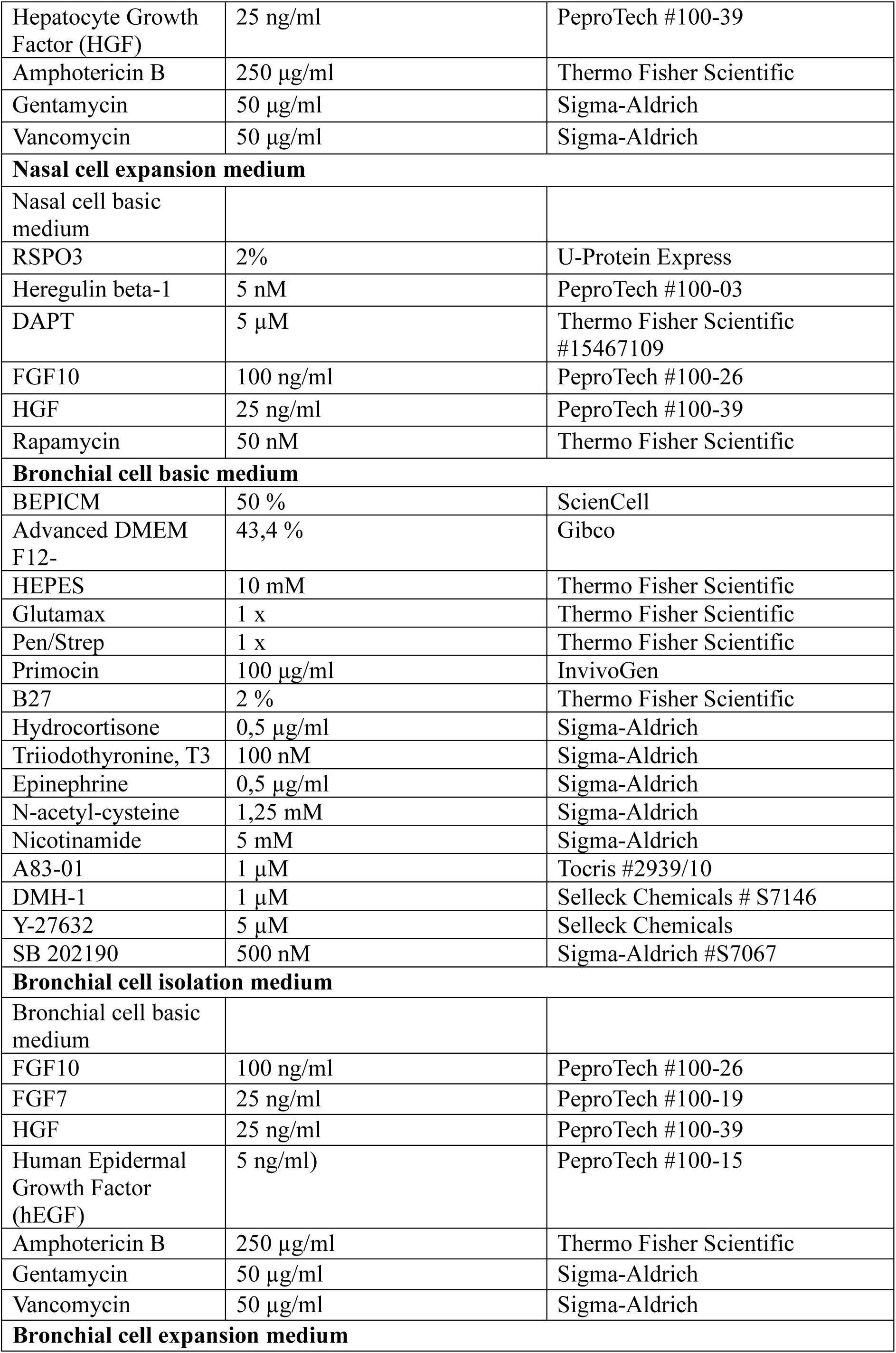
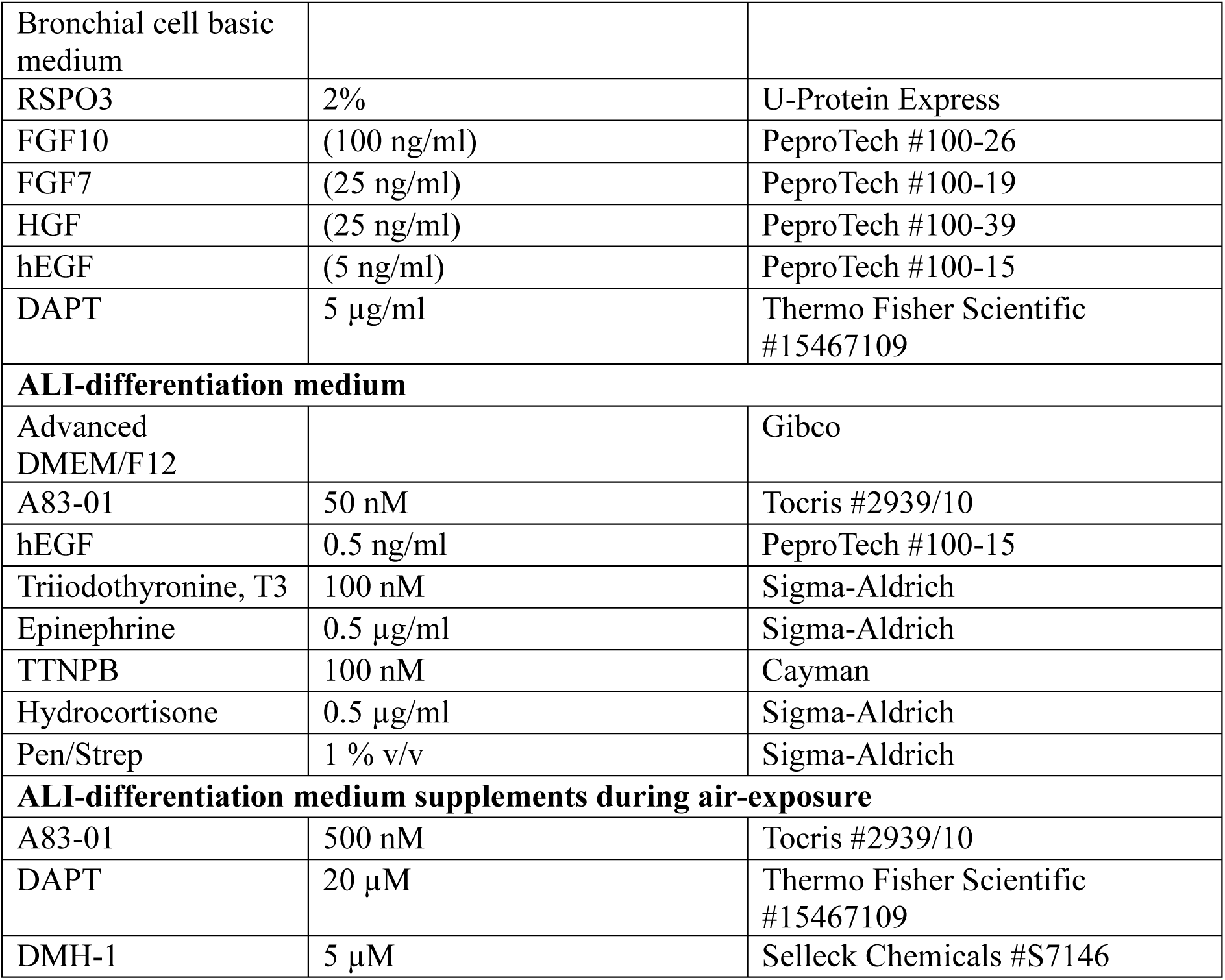
Clinical data and reagents.

## Notes

### Competing Interest Statement

The authors have declared no competing interest.

